# Modelling Intrinsically Disordered Protein Dynamics as Networks of Transient Secondary Structure

**DOI:** 10.1101/377564

**Authors:** Hannah K. Wayment-Steele, Carlos X. Hernández, Vijay S. Pande

## Abstract

Describing the dynamics and conformational landscapes of Intrinsically Disordered Proteins (IDPs) is of paramount importance to understanding their functions. Markov State Models (MSMs) are often used to characterize the dynamics of more structured proteins, but models of IDPs built using conventional MSM modelling protocols can be difficult to interpret due to the inherent nature of IDPs, which exhibit fast transitions between disordered microstates. We propose a new method of determining MSM states from all-atom molecular dynamics simulation data of IDPs by using per-residue secondary structure assignments as input features in a MSM model. Because such secondary structure algorithms use a select set of features for assignment (dihedral angles, contact distances, etc.), they represent a knowledge-based refinement of feature sets used for model-building. This method adds interpretability to IDP conformational landscapes, which are increasingly viewed as composed of transient secondary structure, and allows us to readily use MSM analysis tools in this paradigm. We demonstrate the use of our method with the transcription factor p53 c-terminal domain (p53-CTD), a commonly-studied IDP. We are able to characterize the full secondary structure phase space observed for p53-CTD, and describe characteristics of p53-CTD as a network of transient helical and beta-hairpin structures with different network behaviors in different domains of secondary structure. This analysis provides a novel example of how IDPs can be studied and how researchers might better understand a disordered protein conformational landscape.

---

Tremendous insight has been obtained by viewing proteins as ensembles of metastable structures (1–3), yet the growing study of intrinsically disordered proteins (IDPs) challenges and expands this view (4). IDPs are found ubiquitously in signaling and regulation processes (5), yet their lack of structure poses challenges for studying them using conventional experimental techniques. All-atom molecular dynamics (MD) simulations have the potential to offer atomistic-level detail in the dynamics of IDPs; however, modelling the dynamics of a protein using MD simulations requires describing the conformation space of the protein in a way that lends insight into states and processes of interest to the researcher. Significant study has gone into how to partition this phase space in the field of modelling the dynamics of structured proteins(3), and use of methods for featurization and time-lagged dimensionality reduction (6, 7) have greatly improved the modelling process(8) for proteins that inhabit a few structured conformations. However, in practice, when we use these same tools to analyze MD simulation data of IDPs, we find that these methods face challenges when used to analyze the conformational landscape of an IDP.

## Optimal modelling practices for slow protein dynamics are not necessarily useful for IDPs

There are several aspects of the standard process used to create dynamical models for structure proteins that are less effective for IDPs. To build a MSM using standard protocols (9), all-atom coordinate data is transformed into some set of features, such as the phi/psi backbone dihedrals, closest heavy-atom contact distances, etc. For structured proteins, this space becomes functionally constrained by the finite number of conformations that the protein visits, whereas an IDP that is extended in solution has the possibility to visit many more configurations than a structured protein of the same length.

Next, this featurized data is often decomposed into slow orthogonal dynamical modes using time-structure-based Independent Component Analysis(6, 7, 10) (tICA) or other similar techniques. This allows the modeller to identify slow processes in the data as a function of the input features. Structured proteins typically exhibit only a handful of dynamical modes with long timescales (11, 12). The gap between long timescales and short timescales corresponding to thermal fluctuations is referred to as the “spectral gap”. If only a few slow processes are present, characterizing these processes can be straightforward (Figure 1a). However, when this same modelling protocol is performed on an IDP of interest, we typically observe that many fast processes are identified with little or no spectral gap (Figure 1a). This suggests there are many processes happening at similar timescales, and suggests any time-lagged dimensionality reduction method may be less effective at separating orthogonal modes.

After the data is decomposed into its dynamical processes, it is clustered to identify microstates, and a Markov State Model (MSM) is constructed by counting transitions between these microstates. A MSM provides a description of equilibrium distribution among states and transition rates between them. Pragmatically, structured proteins often occupy just a limited number of states of interest when compared to IDPs (Figure 1b). For instance, a kinase may have a few active/inactive states or a protein folding may have only a handful of folding intermediate states of interest. In contrast, models of IDPs may exhibit hundreds of microstates that all appear similarly disordered to the researcher’s eye, and can be difficult to distinguish or interpret.

**Figure 1:**
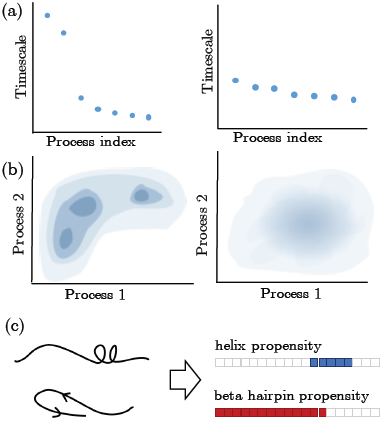
Challenges of modelling IDPs with traditional methods for modelling dynamics of structured proteins (a): structured proteins typically exhibit a few slow dynamical modes. On the other hand, IDPs exhibit many fast modes with no spectral gap. (b) Structured protein conformational landscapes are typically characterized by a finite number of basins of interest. Conversely, IDPs inhabit many varied microstates that may all appear disordered to the modeller without further tools for interpretation. The interpretation tool presented here (c) is to compute the secondary structure assignments of MD simulation data of the IDP in order to view the conformational landscape in terms of its transient secondary structure, in line with current experimental techniques.

The above limitations of conventional structured protein analysis methods on IDPs follow researchers’ intuition about IDPs, that they are rapidly interconverting between transient states and therefore may not exhibit slow dynamical processes. Given these characteristics of IDPs, how can we effectively model IDPs in an interpretive way? Any modelling process must balance a tradeoff between interpretability and accuracy. The above methods for structured proteins can be optimized for accuracy in their representation of slow dynamical modes (12, 13), but perhaps to make headway into understanding IDPs, we need to take a step back and think about how we might best interpret disordered protein landscapes, and how we can create models to further our aims of interpretation.

## Secondary structure assignment projects IDP conformations to a more interpretable paradigm

To attempt to create IDP conformational landscape models that are more intuitive, we drew inspiration from experimental fields that have had success in thinking about IDPs in terms of secondary structure assignment (14). For instance, the *δ*2D method is able to determine per-residue secondary structure populations directly from chemical shifts (15). Work such as this has contributed to a paradigm in which IDPs exhibit fast transitions between transient secondary structure conformations, some of which then are stabilized upon binding to target proteins. However, experiments are limited when a measurement, such as secondary-structure propensity, is only able to represent an ensemble average over microstates (14). This limitation can be addressed by complementing experiment with molecular simulation, which offers atomic-level resolution of an ensemble of protein structures. In this light, we propose using this transient-secondary-structure paradigm of thinking to parse MD simulation data prior to building a MSM with it. We use the output of a secondary structure assignment algorithm as a featurization for MD simulation data of an IDP. This method allows us to fully partition and characterize the phase space of the IDP as thought of in terms of its transient secondary structure.

## Secondary structure assignments reveal a kinetic network of transient structure

To demonstrate this method, we analyze the commonly-studied IDP, the C-terminal domain of the p53 transcription factor (p53-CTD). This peptide was simulated for a total time of over 6 ms on the distributed computing platform Folding@Home(16) in the force field Amber ff99SB-ILDN (17). We computed secondary structure assignments for each residue at each time point with the commonly-used secondary structure assignment tool DSSP (18). For the purpose of demonstration here, we use the simplified assignment set (helix, beta strand, or random coil), but note that the same analysis would be possible with the more complete set of 8 possible assignments, or with other secondary structure assignment tools entirely. Because secondary structure assignment is a categorical variable, we encode each assignment per residue as two dummy variables representing the state of having helical or beta-hairpin character, i.e. (1,0) if assigned as helical, (0,1) if beta strand, or (0,0) if random coil.^1^ We then cluster the data using the minibatch K-Means algorithm and use the cluster assignments to construct a MSM.

Viewing the landscape of p53-CTD as a network immediately offers several immediate observations (Figure 2). In a network representation of a MSM for p53-CTD featurized using the DSSP secondary structure prediction, we represent microstates as nodes. Microstates are positioned according to their connectivity degree, so nodes closer to the center are more hub-like while the nodes on the periphery are less connected. The weights of edges between nodes represent the interconversion rates between nodes.

**Figure 2:**
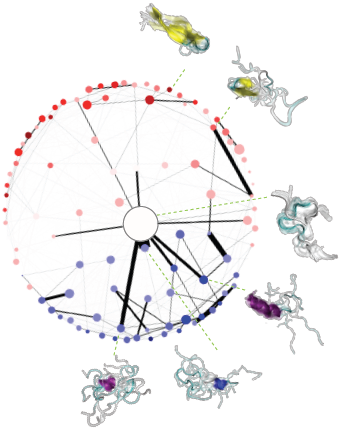
By visualizing p53-CTD conformational landscape as a network, we can immediately gain insight into its network properties. The entirely disordered state acts as a kinetic hub for a landscape with a majority of microstates holding transient beta-hairpin character and a minority with helical character. However, several helical microstates have prominent hub-like character as well. Several prominent microstates are depicted surrounding the network as examples.

The microstate with the predominant population (roughly 24%) is that with no secondary structure assignment. The converse of this is that 76% of the conformational space can be described by some degree of secondary structure assignment, which suggests that thinking of IDP landscapes using the secondary structure paradigm is useful. From the network visualization, we can observe that the disordered state serves as a kinetic hub for p53-CTD to interconvert between states with more secondary structure. This observation is perhaps intuitive, yet viewing the conformational landscape as a network analysis allows us to use relevant network analysis tools to draw conclusions. We furthermore observe that p53-CTD has more transient states with beta-structure propensity than with helical propensity; yet the helical microstates exhibit more interconversion.

**Figure 3:**
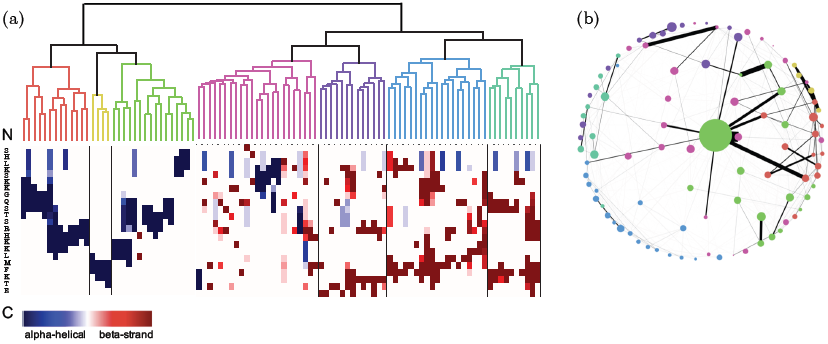
Hierarchically clustering the microstates (a) with MVCA allows us to observe which states are kinetically similar. In (b), we color the same network model by clustered microstate assignment for comparison to Figure 2, observing that kinetically similar microstates are also close to one another in the network.

We can also apply other current MSM analysis tools to interpret the p53-CTD landscape. We use hierarchical clustering of microstates (MVCA)(19) to understand which microstates are kinetically close to one another. The resulting analysis allows us to better visualize the span of observed transient secondary structure. For instance, we observe that there are a limited number of locations along the IDP where helices and beta-strand motifs can form.

## Suitability of Timescale-optimized models

A pertinent question is how this secondary-structure-based featurization compares to other featurizations in existing measures of model quality. For MD data analysis, model quality is often evaluated in terms of the Variational Approach to Conformational Dynamics, which states that no process may be detected in data that is slower than the true slowest process. This principle implies that a model with longer characteristic timescales is closer to representing the true processes in the data than a model with shorter characteristic timescales. This principle has been used to develop several methods for maximizing eigenvalue-based scores (12, 20, 21), one of these being the Generalized Matrix Rayleigh Quotient (GMRQ) (12).

We compare the GMRQ scores of models built on p53-CTD with three different featurizations: secondary structure (the method presented in this work), *ϕ*/*ψ* backbone dihedral angles, and closest heavy-atom contacts between residues. Backbone dihedral angles and contacts are frequently used for featurization and have been shown to provide high GMRQ scores for structured proteins (8). We note that secondary-structure prediction algorithms classify structures based on a function of these other features (18). By featurizing with secondary structure assignment, we are selecting features previously identified as pertinent to defining secondary structure phase space through expert knowledge. In the context of structured proteins, where a helix in one conformation will likely remain a helix across conformations, this is perhaps not so useful, but for IDPs, which are rapidly switching between different secondary structure states, this simplifies the scope of possible feature space to characterize.

Because secondary structure creates a more coarse-grained phase space than either dihedrals or contacts, we expect secondary structure models to not perform as well in optimizing timescales as a sacrifice in the name of interpretability. We observe that DSSP featurization does perform more poorly than models built with dihedrals, (i.e. is unable to find as long of timescales), but performs comparably to models built with contacts (Figure 4a). Parameter optimization with Osprey was performed using dihedrals, contacts, or secondary structure as input, varying the lag time and number of components used for tICA, and the number of clusters used for clustering across a range of values.

**Figure 4:**
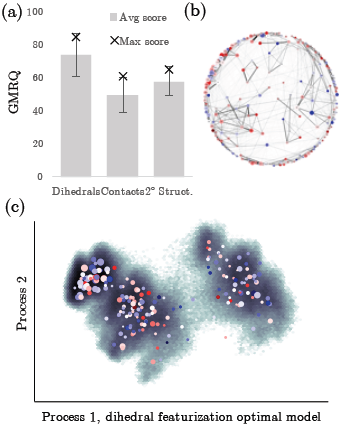
Dihedral angles produce the highest average GMRQ score, a measure of model quality in representing slow timescales, yet are less interpretable in terms of secondary structure. Albeit being a much simpler representation, the secondary structure featurization performs slightly better than contact distances, another commonly-used featurization, although does not capture as slow of timescales as dihedral angles. (a). Error bars represent one standard deviation. A variationally optimal model featurized by dihedral angles is less interpretable in terms of transient secondary structure (b,c). When the optimal dihedral model is viewed as a network (b), microstate secondary structure is uncorrelated from the network structure, unlike in the network from the MSM model featurized by secondary structure. Likewise, mapping data onto two slowest processes identified does not map to interpretable secondary structure trends. Microstates are sized by their populations and colored by the computed secondary structure propensity of samples drawn from each microstate.

Although dihedral featurization may give higher GMRQ scores, models built from these are more difficult to interpret as a landscape of secondary structure. We find that the model with the highest GMRQ score does not identify any interpretable trends in secondary structure propensity either when viewed as a network model (Figure 4b) or when projected onto the slow processes identified (Figure 4c). While using a GMRQ-optimized model might certainly be preferable for analyzing other observables, such a model may be much more difficult to use such a model to understand an IDP as a collection of microstates with varying secondary structure.

In summary, we introduce a new method for featurizing MD trajectory data of IDPs that draws inspiration from viewing IDPs in terms of transient secondary structure assignments. We demonstrate that it creates readily interpretable network models and discuss the philosophical tradeoff between interpretability and objective model quality. We hope the use of such tools will improve the study and analysis of IDPs and continue discussion of how we may best characterize disordered conformational landscapes.

## AUTHOR CONTRIBUTIONS

HKWS, CXH, and VSP conceived of the research. CXH performed the MD simulations. HKWS designed and performed the analysis. HKWS and CXH wrote the manuscript.

## ACKNOWLEDGMENTS

The authors thank B. E. Husic and M. M. Sultan for useful and insightful discussion. HKWS and CXH acknowledge support from NSF GRFP (DGE-114747). We graciously acknowledge the Folding@home donors who contributed to the p53-CTD simulations. VSP is a consultant and SAB member of Schrodinger, LLC and Globavir, sits on the Board of Directors of Apeel Inc, Freenome Inc, Omada Health, Patient Ping, Rigetti Computing, and is a General Partner at Andreessen Horowitz. Wayment-Steele, Hernández, Pande

## Footnotes

1 While we could use three variables to represent the three possible assignments (helix, beta strand, or random coil) in a one-hot encoding style, this is overrepresented and we choose not to for the sake of minimizing the total amount of data to be processed.

